# Cell Proliferation and Morphogenetic Compartmentalization in the Phoronid *Phoronopsis harmeri*: Conserved and Derived Patterns

**DOI:** 10.1101/2025.09.19.677378

**Authors:** Evgeny G. Ivashkin, Olga I. Taimanova, Anton I. Bogomolov, Elena N. Temereva

## Abstract

Cell proliferation is a key driver of morphogenesis and body plan transformation in multicellular animals, yet its spatial organization remains poorly understood in many non-segmented spiralians. In this study, we examine the dynamics of cell division during larval growth and metamorphosis in the larvae and early juveniles of the phoronid *Phoronopsis harmeri*, using EdU incorporation, anti-phospho-histone H3 immunostaining, confocal laser scanning microscopy, and electron microscopy. Early larval development is characterized by widespread proliferative activity across ectodermal and mesodermal tissues, which becomes progressively compartmentalized as the larva matures. Two structured ring-shaped posterior proliferative zones, pre- and post-telotrochal, emerge within the telotroch and persist through metamorphosis, supporting both larval elongation and the juvenile development of ascending gut branch. In contrast, the metasomal sac and future adult trunk epidermis expand via broadly distributed epithelial proliferation, without forming a localized growth zone. This suggests that *P. harmeri* combines conserved features, such as a posterior growth zone, with lineage-specific innovations in regional growth. In addition, we identify atypical mitotic characteristics in this species, including unconventional metaphase organization and signs of interkinetic nuclear migration in larval epithelia. Our findings highlight the coexistence of ancestral and derived proliferative mechanisms in phoronids and provide new insights into the evolution of axial elongation and morphogenetic compartmentalization in Lophotrochozoa.

## Introduction

Growth zones in developing embryos of multicellular animals are regions with an increased number of proliferating cells compared to surrounding tissues. This localized proliferation results from the activity of specific morphogen gradients that regulate tissue patterning during organogenesis (Rogers and Schier, 2011). These zones play a crucial role in coordinating of body elongation, segmentation (where present), and organ differentiation. Such proliferation zones either can be conserved ancestral features or may have been evolved secondarily as part of body plan modifications in response to ecological and functional demands. In some groups, growth zones are maintained throughout development, ensuring progressive segment addition, as observed in annelids (Balavoine, 2014). In others, they appear transiently or shift in function depending on developmental stage (Seaver et al., 2005; Steventon et al., 2016). Additionally, evolutionary trends toward miniaturization or simplification of body plans may lead to the loss or reorganization of growth zones, as seen in certain parasitic and sessile organisms.

Spiralia, one of the major clades of bilaterian animals, exhibits remarkable diversity in developmental strategies and body organization. This superphylum includes annelids, mollusks, phoronids, nemerteans, bryozoans, and platyhelminths, among others. While annelids are characterized by a segmented body plan, many other lophotrochozoans exhibit non-segmented body structures, raising important evolutionary and developmental questions regarding the role of cell division patterns and the presence of a posterior growth zone (PGZ) in axis elongation. Understanding these patterns in both segmented and non-segmented lophotrochozoans provides insight into the evolution of metazoan development and segmentation (Rogers and Schier, 2011).

Phoronids, brachiopods, and bryozoans, three groups linked due to their lophophore-bearing morphology, also display distinct growth patterns (Kuzmina and Temereva, 2024). However, information regarding the organization of proliferation zones in these animals remains extremely limited. In bryozoans, cell proliferation contributes to continuous budding and colony growth rather than posterior elongation (Borg, 1926; Fuchs et al., 2011). It has also been observed that cells incorporating EdU are localized at the base of tentacles during their development. Additionally, tentacle growth is associated with cell divisions at its base during regeneration (Shunatova and Borisenko, 2020). In phoronids, proliferative activity was initially described only briefly (Bird, 2012), with subsequent studies focusing primarily on the tentacles of larvae and metamorphic individuals (Temereva, 2025). In brachiopods, data on experiments with visualization of proliferative activity have never been published. Further research on these animal groups could provide crucial insights into the evolution of growth and body patterning, not only within Lophotrochozoa but also across Bilateria as a whole.

Phoronid development includes several crucial stages: from embryogenesis through the early bilobed larva, followed by extended larval development and culminating in catastrophic metamorphosis (Temereva and Kostyuchenko, 2025; Temereva and Malakhov, 2015, 2012, 2007; Temereva and Tsitrin, 2014, 2014; Zimmer, 1964). After fertilization, the blastula forms approximately 16 hours after polar body formation and persists for several hours as cells continue to proliferate and differentiate. Gastrulation is completed within the first 24 hours post-fertilization, resulting in the formation of embryonic germ layers and the establishment of the basic body plan (Temereva and Malakhov, 2007). Larval development is characterized by progressive body growth and tentacle elongation (Temereva and Neretina, 2013). Metamorphosis in phoronids represents a profound reorganization of multiple larval organ systems and a comprehensive transformation of the body plan (Herrmann, 1976; Temereva, 2010; Temereva and Malakhov, 2015; Temereva and Tsitrin, 2014).

In the present study, we investigated the distribution of cell divisions during larval growth and metamorphosis in *Phoronopsis harmeri* using EdU labeling, immunocytochemistry, and confocal microscopy. Our findings indicate that a posterior growth zone is established and functions within the epithelium (specifically in telotroch) and the gut of the actinotroch larva, persisting until metamorphosis. Simultaneously, the growth of the metasomal sac and the subsequent expansion of the post-metamorphic body covering occurs through scattered cell divisions rather than a distinct localized growth zone. This suggests that the growth of adult body coverings represents an evolutionarily novel trait, whereas the posterior growth zone in the actinotroch larva reflects a conserved developmental mechanism, reinforcing its identity as a secondary larval stage.

## Experimental Procedures

### Animal сollection and fixation

Larvae of *Phoronopsis harmeri* Pixell, 1912 were collected in Vostok Bay (Sea of Japan) using a plankton net during November 2022 and 2024. The specimens were cultured in natural seawater at 13–14□°C under controlled laboratory conditions until metamorphosis. Prior to fixation, animals were anesthetized and relaxed in a 7% magnesium chloride (MgCl₂) solution prepared in filtered seawater. Individuals at different developmental stages were then processed for analysis by transmission electron microscopy (TEM), scanning electron microscopy (SEM), EdU incorporation assay, and immunohistochemistry (IHC).

### Scanning (SEM) and transmission (TEM) electron microscopy

For SEM and TEM, specimens were fixed in 2.5% glutaraldehyde in 0.1□M cacodylate buffer and washed in the same buffer for 12 hours, following protocols described previously (Temereva and Malakhov, 2007; Temereva and Tsitrin, 2014).

For SEM, fixed specimens were dehydrated through a graded ethanol-acetone series, critically point dried, and sputter-coated with a platinum–palladium alloy. Imaging was performed using a JEOL JSM-6380 scanning electron microscope.

For TEM, the specimens were post-fixed in 1% osmium tetroxide in phosphate-buffered saline (PBS) for 2 hours, washed in PBS, dehydrated through an ethanol-isopropyl alcohol series, and embedded in Embed-812 resin. Semithin and ultrathin sections were prepared with a Leica EM UC-7 ultramicrotome. Semithin sections were stained with methylene blue and examined under an Olympus BX51 microscope equipped with an AxioCam HRm camera. Ultrathin sections were stained with uranyl acetate followed by lead citrate and analyzed using a JEOL JEM-1011 transmission electron microscope.

### EdU cell proliferation assay and “сlick” сhemistry

To assess cell proliferation, *P. harmeri* larvae at various developmental stages were incubated with 200□μM EdU diluted in filtered seawater for 5 or 45 minutes. Specimens were then washed with filtered seawater and with 7% MgCl₂ in seawater, followed by fixation in 4% paraformaldehyde in 1.75× PBS with 0.1% Tween-20 (PTw) for 3 hours at room temperature. After fixation, larvae were washed in PTw, dehydrated, and stored in 100% methanol at −20□°C.

To remove fixative, larvae were rehydrated and washed three times for 10 minutes in 0.1% Triton X-100 in 0.01□M PBS (pH 7.4) at room temperature. They were then incubated in blocking solution (3% BSA, 0.1% Triton X-100 in 0.01□M PBS) for 1 hour. Following this, samples were washed three times for 10 minutes in Tris-NaCl reaction buffer (50□mM Tris, 150□mM NaCl, pH□7.5) and transferred into the Click-iT reaction cocktail prepared according to the manufacturer’s protocol (Vector Laboratories, EdU Cell Proliferation Assay Protocol for Fluorescent Microscopy).

The reaction cocktail consisted of 445□μL Tris-NaCl buffer, 5□μL 100□mM CuSO₄, 1.2□μL Alexa Fluor® 647 azide (Molecular Probes), and 50□μL 55□mM sodium ascorbate (Sigma-Aldrich). Larvae were incubated in the cocktail for 1 hour, then washed once in reaction buffer (5□min), once in blocking buffer (10□min), and finally twice in washing buffer (PBS 0.01□M with 2□mM NaN₃ and 0.5□mM EDTA) followed by a final 15-minute rinse in 0.01□M PBS.

### Immunohistochemistry

Immunodetection of phospho-histone H3 was performed after the Click reaction. Larvae were incubated with anti-phospho-histone H3 (Ser28) rat monoclonal antibody (Sigma-Aldrich, H9908) diluted 1:1000 in blocking solution, for 24 hours at 4□°C. After incubation, samples were washed three times for 15 minutes in 0.1% Triton X-100 in 0.01□M PBS.

Secondary detection was carried out using goat anti-rat IgG conjugated with Alexa Fluor® 546 (Molecular Probes), diluted 1:800, for 12 hours at 4□°C. Following incubation, larvae were washed three times for 5 minutes each in 0.01□M PBS.

### Confocal laser scanning microscopy (CLSM), image processing, and morphometric measurements

Fluorescent imaging was performed using a Zeiss LSM 880 laser scanning confocal microscope. Image processing, volume rendering, and three-dimensional visualization were conducted in Imaris 9.9 (Bitplane, Oxford Instruments). Morphometric measurements were carried out directly in Imaris on confocal image stacks of whole-mount specimens labeled with EdU and DAPI.

To assess changes in larval body proportions during development, we measured body length from the mouth to the anterior edge of the telotroch, preoral lobe width, and telotroch diameter based on confocal image stacks, following the scheme shown in Fig.□1H1. Measurements were analyzed in GraphPad Prism 9 to generate plots and calculate mean values ± standard error of mean (SEM). Larvae selected for measurement were staged according to external morphological criteria, including tentacle number, telotroch width, and overall body shape. At least five larvae were analyzed per developmental stage (early, mid, advanced, and competent).

## Results

### Developmental stages, growth, and timing of *Phoronopsis harmeri*

The external morphology of *P. harmeri* larvae changes significantly throughout development (Fig. 1A-D). At early stages, the larva possesses a small preoral lobe at the anterior end, a developing collar region with shallow tentacle buds, and an elongated, slender trunk. As the larva grows, the preoral lobe enlarges and becomes more dome-shaped, while the tentacles elongate and increase in number, forming a prominent crown encircling the mouth. The trunk gradually thickens and elongates, and at the ten-tentacle stage, the metasomal sac, initially a minor ventral depression, deepens into a pouch-like structure. During mid-larval development, the telotroch, a ciliated band at the posterior end, becomes more pronounced and widens, aiding in effective swimming. In competent larvae, the body is distinctly divided into three regions: a broad, rounded preoral lobe bearing the apical organ, a wide collar bearing 24 elongated ciliated tentacles and containing spacious blastocoel where the blood masses are located, and a robust trunk with a visible metasomal sac on the ventral side and a large, circular telotroch at the posterior. These external modifications reflect the larva’s preparation for settlement and metamorphosis into the benthic juvenile form.

**Figure 1.**
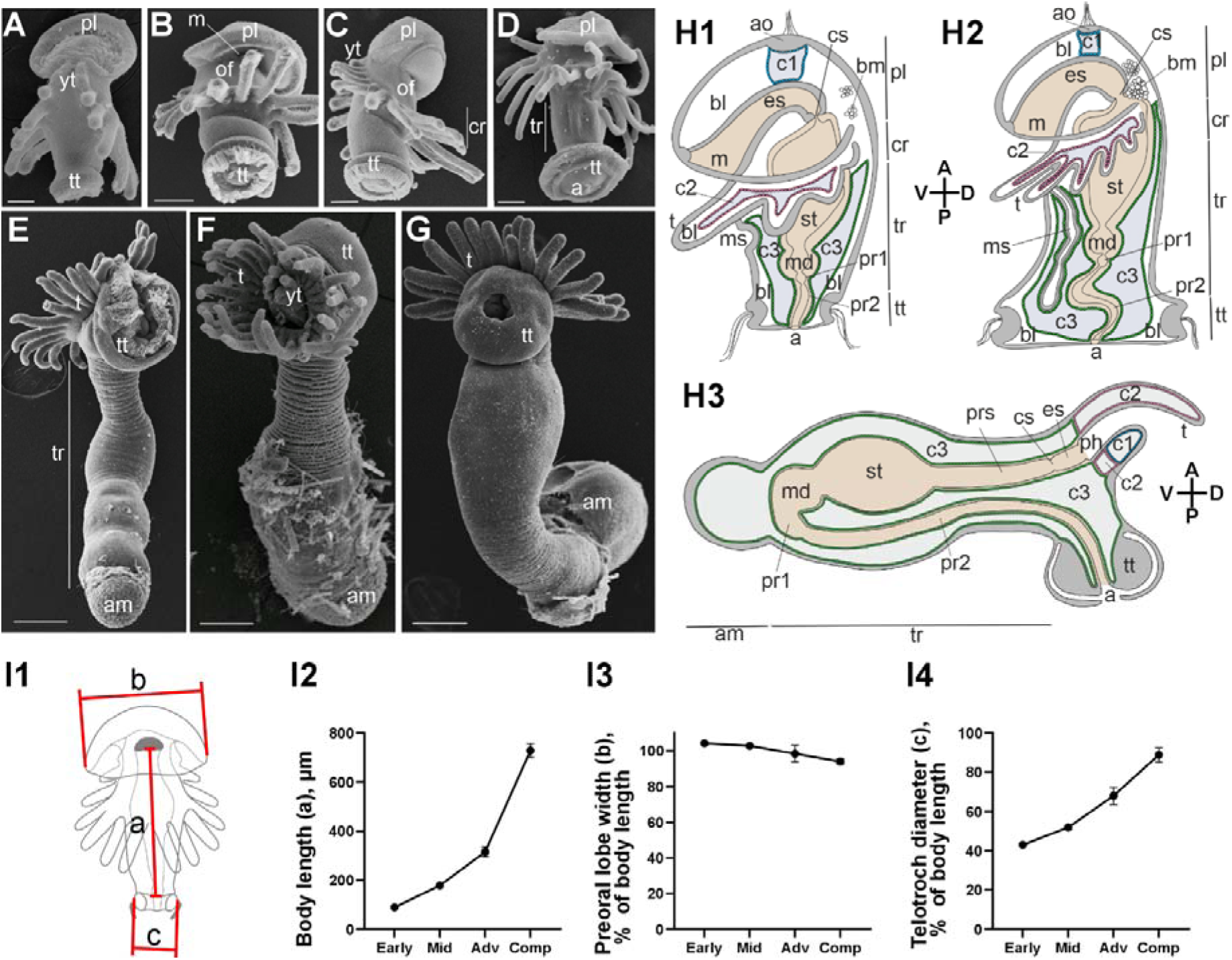
External morphology and growth dynamics of *Phoronopsis harmeri* at different stages of development. (A–G) Scanning electron micrographs (SEM) of larvae and juveniles: (A) 10-tentacle larva; (B) 12-tentacle larva; (C) 20-tentacle larva; (D) competent larva; (E) early metamorphic juvenile (posterior view); (F) mid metamorphic juvenile (anterior view); (G) mid metamorphic juvenile (posterior view). (H1–H3) Schematic drawings of body organization: (I1) 10-tentacle larva; (I2) competent larva; (I3) metamorphic juvenile. (I1–I4) Growth dynamics during larval development (mean ± SEM, N = 5): (H1) scheme of measurements; (H2) mouth to telotroch body length (a on the scheme, μm); (H2) preoral lobe width as a percentage of body length (b on the scheme); (H3) telotroch diameter as a percentage of body length (c on the scheme). Note that the preoral lobe grows proportionally with overall larval size, while the telotroch increases disproportionately, becoming significantly larger relative to the trunk. Abbreviations: A – anterior, a – anus, am – ampulla, ao - apical organ, bl – blastocoel, bm - blood mass, c1 – protocoel, c2 – mesocoel, c3 – metacoel, cr – collar region, cs - cardial sphincter, D – dorsal, es – esophagus, m – mouth, md – midgut, ms - metasomal sac, of - oral field, P – posterior, ph – pharynx, pl - preoral lobe, pr – proctodaeum, pr1 – proctodaeum: part one, pr2 – proctodaeum: part two, prs – prestomach, st – stomach, t – tentacle, tr – trunk, tt – telotroch, V – ventral, yt - youngest tentacle. Scale bars: A – 50 µm; B–D – 100 µm; E–G – 200 µm.

Larval development progresses slowly. It takes about 24 days post-fertilization for larvae to reach the six-tentacle stage, which represents early larval growth. The transition from the six-tentacle to the eight-tentacle stage occurs gradually over several days, with continuous tentacle addition observed. The period between the eight-tentacle and twelve-tentacle stages also spans several days to a week, reflecting a steady but slow morphogenesis. Development from the twelve-tentacle to the twenty-tentacle stage can take an additional one to two weeks, depending on environmental conditions such as temperature and food availability.

The attainment of twenty-four tentacles corresponds to larval competence for metamorphosis, typically occurring three months post-fertilization. The initial stage of metamorphosis, once initiated, is a relatively rapid process, with all catastrophic changes occurring within approximately half an hour (Fig. 1E-G). It includes the eversion of the metasomal sac and the relocation of the gut into it, along with the contraction of the larval body wall and the telotroch. This stage involves both the resorption of larval structures – such as the preoral lobe with the apical organ and preoral ciliated band, the postoral ciliated band, and the telotroch – and the concurrent transformation of the everted metasomal sac into the adult body.

In *P. harmeri*, before metmorphosis, the organization of the coelom is characterized by a tripartite pattern: the protocoel is located in the preoral lobe under the apical organ, the mesocoel resides at the bases of the tentacles, and the metacoel fills most of the trunk region, separated from the body wall by a spacious blastocoel (Fig. 1H1-H2). Upon the initiation of the metamorphosis, the metasomal sac everts and induces profound reorganization of body cavities (Fig. 1H3).

The morphogenesis of the digestive tract in *P. harmeri* undergoes a series of profound transformations from the early larval stages through metamorphosis to the juvenile form (Fig. 1H1-H3). In actinotroch larvae, the gut is composed of a straight digestive tube consisting of an ectodermal esophagus, a bilobed stomach (upper and lower chambers), an endodermal midgut, and an ectodermal proctodeum. As the larva approaches competence, the proctodeum becomes highly elongated and forms several loops within the trunk coelom. During the initial stage of metamorphosis, the metasomal sac everts, pulling the digestive tract into a newly forming ventral body region and giving rise to the characteristic U-shaped gut of the juvenile. This rearrangement is accompanied by straightening of intestinal loops and the reconfiguration of the digestive tube into descending and ascending branches. Subsequently, distinct morphological regions within the hindgut become discernible, including the progressive formation of a terminal anal chamber from an ectodermal invagination.

To characterize the dynamics of growth and changes in body proportions during larval development of *P. harmeri*, we measured the body length (from the mouth to the upper border of the telotroch), the width of the preoral lobe, and the diameter of the telotroch (Fig. 1I1). We estimated that over the course of larval development, the body length increases by approximately eight-fold (Fig. 1I2). Notably, the proportions of the larval body also undergo significant modifications during growth. While the preoral lobe expands proportionally relative to the overall body size, maintaining its shape and relative dimensions (Fig. 1I3), the telotroch exhibits a disproportionate increase in size, growing approximately two times relative to the initial proportion observed in early larvae (Fig. 1I4).

Overall, *P. harmeri* displays a protracted larval development period, with significant durations at each developmental stage, reflecting its adaptation to planktonic life and the transition from pelagic to benthic life.

### Features of mitosis in *Phoronopsis harmeri*

To investigate the nature of cell divisions in phoronids, we employed a combination of transmission electron microscopy (TEM), EdU incorporation assays, and phospho-histone H3 (pH3) immunostaining.

TEM analysis revealed that metaphase plates in *P. harmeri* exhibit a highly atypical organization compared to the canonical arrangement in most metazoans (Fig. 2A1–A5). Chromosomes are not aligned along a defined equatorial plane but instead form compact, irregularly shaped clusters dispersed throughout the cytoplasm. These chromosomal aggregates occupy a large portion of the intracellular space, often displacing or entirely replacing organelle-free regions of the cytoplasm. Although chromosomes appear maximally condensed, their morphological resolution is surprisingly poor: sister chromatids are not clearly separated, centromeric or kinetochore structures are indistinct or absent, and chromosome contours are smooth and relatively homogeneous. Furthermore, the electron density of metaphase chromatin (Fig. 2A2-A5) is not elevated relative to surrounding structures – in fact, it is sometimes lower than that of interphase heterochromatin (Fig. 2A1), suggesting an unconventional mode of chromatin condensation. The internal definition of metaphase chromosomes is inferior to that of the dense heterochromatin clusters observed in interphase nuclei, where euchromatin–heterochromatin boundaries are clearly demarcated. Across all mitotic stages, no well-organized or prominent mitotic spindles were visible by TEM.

**Figure 2.**
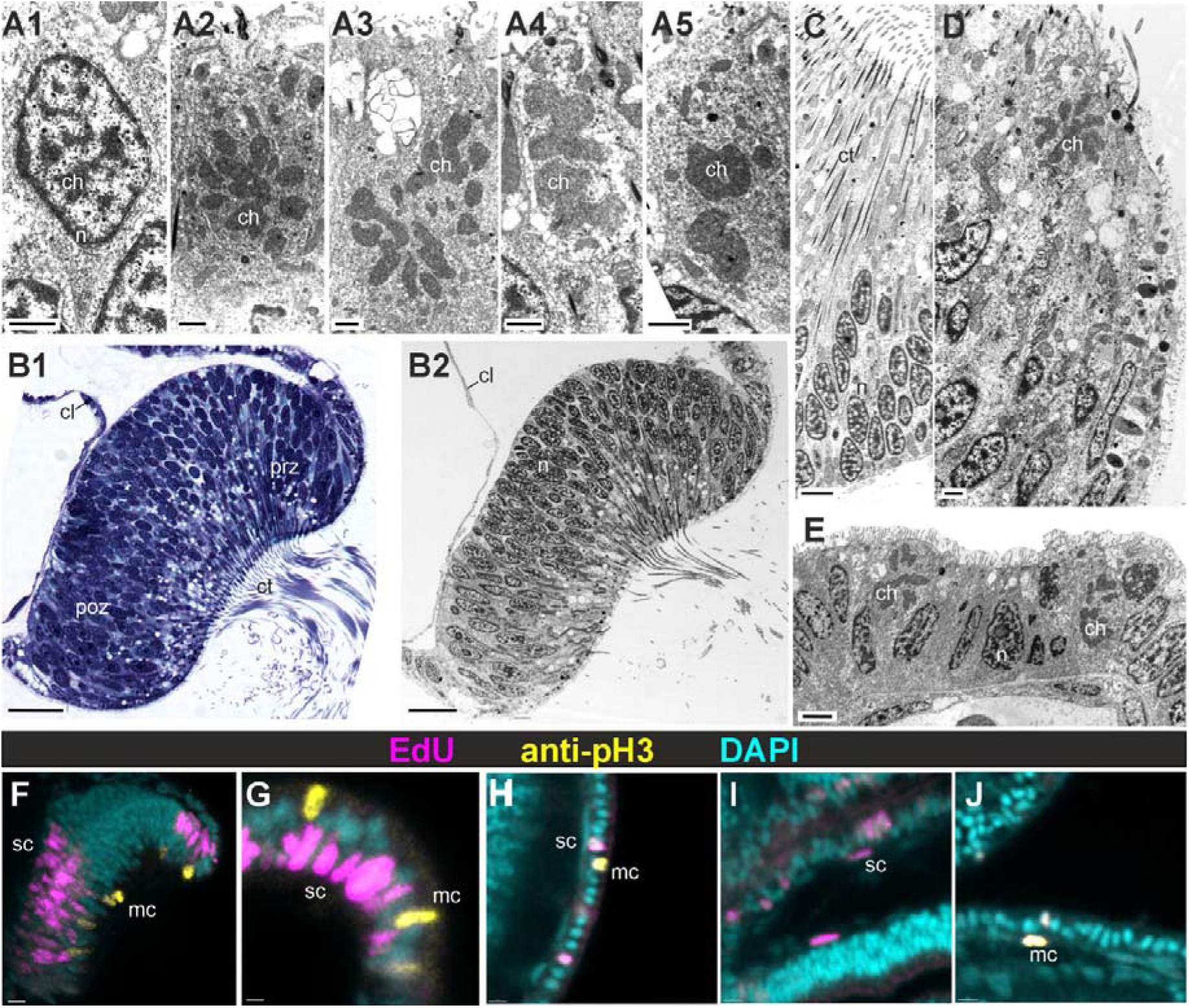
Characteristics of cell division in larvae of *Phoronopsis harmeri*. (A1–A5) Nuclei at different stages of the cell cycle. (B1, B2) Transverse sections through the telotroch ring. (C, D, E) Enlarged views of the telotroch epithelium: (C) central zone with slender cells bearing long cilium; (D) peripheral pre-telotrochal proliferation zone, where mitotic cells are concentrated; (E) peripheral post-telotrochal proliferation zone. (F–J) EdU⁺ and pH3⁺ cells in various larval regions: (F) transverse section of the telotroch ring; (G) epithelium near the mouth; (H) epithelium of the trunk at the mid-body level; (I, J) proliferating cells in the lining of the trunk coelom. (A1-A5; B2; C-E) TEM; (B2) semithin section stained with toluidine blue; (F-J) CLSM. Magenta – EdU-click; yellow – anti-phosphorylated histone H3 IHC; cyan – nuclei stained with DAPI. Abbreviations: ct – central ciliated zone of telotroch, cl – coelomic lining, ch – chromosome, mc - M-phase cells, n – nucleus, poz - post-telotroch proliferation zones, prz - pre-telotroch proliferation zones, sc – S-phase cells. Scale bars: A1–A5, D – 1 µm; C – 3 µm; E – 2 µm; B1–B2 – 10 µm; F–J – 5 µm.

In the telotroch, which possesses a pseudostratified epithelium composed of monociliated cells, cell divisions exhibit a consistent orientation perpendicular to the epithelial surface (Fig. 2B1–D). During prometaphase, coinciding with nuclear envelope breakdown, the nuclear contents migrate toward the apical (outer) surface of the epithelium (Fig. 2D). Apical protrusion of metaphase nuclei is also observed in other epithelial regions of the larval body (Fig. 2E).

EdU-positive (EdU⁺) nuclei, indicative of S-phase cells, are predominantly localized near the basal surface, while phospho-histone H3-positive (pH3⁺) mitotic figures accumulate apically (Fig. 2F). Notably, the spatial separation of EdU⁺ and pH3⁺ nuclei is not unique to the telotroch; elevated metaphase nuclei are similarly found in monolayered surface epithelia throughout the larval body (Fig. 2G, H).

In contrast to the telotroch and epidermis, the coelomic lining displays mitotic spindles oriented parallel to the epithelial plane and lacks signs of nuclear displacement. No apicobasal segregation of EdU⁺ and pH3⁺ nuclei is observed in these mesodermal cells, suggesting that interkinetic nuclear migration is absent in the coelomic epithelium (Fig. 2I, J). Additionally, in both the surface epithelia and coelomic lining, pH3 labeling does not form compact metaphase chromosomes but instead appears as a diffuse signal distributed across a substantial portion of the cell volume (Fig. 2F–J).

Together, these findings suggest that the process of mitosis in phoronids involves species-specific modifications at both tissue and subcellular levels. While the chromosomal architecture during cell division displays low contrast and spatial irregularity, these characteristics do not hinder the use of standard methods for analyzing cell proliferation during the development of *P. harmeri*.

### Spatial patterns of cell proliferation in larval stages and metamorphic juveniles of *Phoronopsis harmeri*

The earliest stage analyzed in our study was the 10-tentacle actinotroch (Fig.3A1-A3). At this point, the telotroch is nearly formed, and the invagination of the metasomal sac has just begun. EdU⁺ and pH3⁺ cells are widespread across the epidermis and within the coelothelium of all coelomic compartments (Fig. 3A1–A3, B, D). Both the dorsal and ventral surfaces of the preoral lobe show active proliferation, as does the developing esophagus (Fig. 3B). Within the preoral lobe, elevated mitotic activity is observed in the region corresponding to the apical organ. Intensive proliferation is particularly prominent at the bases of the tentacles (Fig. 3C). In contrast, only a few EdU⁺ or mitotic cells are found in the stomach walls (Fig. 3B, D), although these areas display nonspecific cytoplasmic fluorescence. At the same time, numerous dividing cells are present in the midgut and proctodeum.

**Figure 3.**
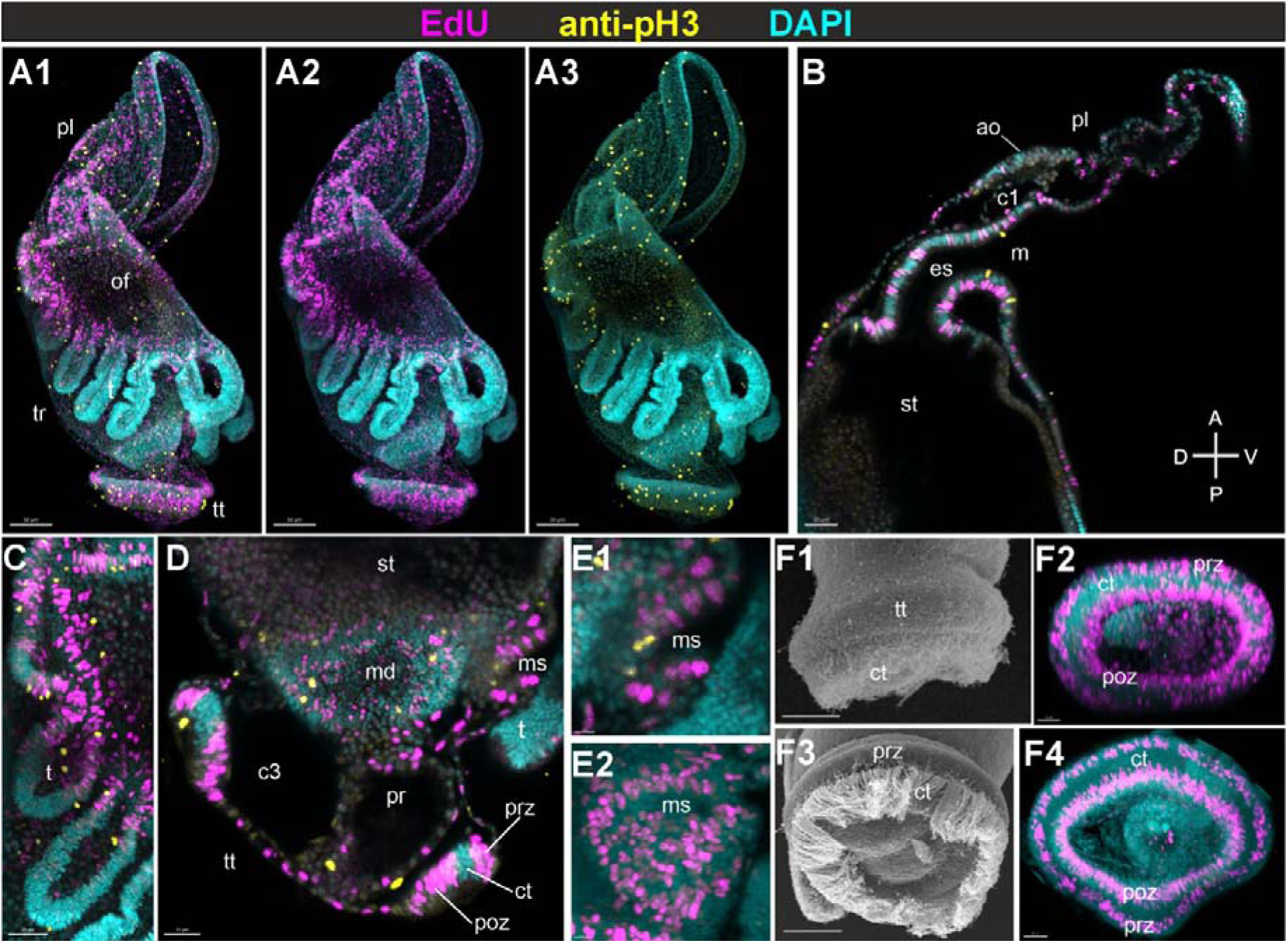
Proliferative activity in early-stage larvae of *Phoronopsis harmeri*. (A1–A3) Whole 10-tentacle larva, lateral view. (B, C) Sagittal optical sections through anterior (B) and posterior (C) body regions. (D1, D2) Metasomal sac: (D1) sagittal section (10-tentacle larva); (D2) frontal section (12-tentacle larva). (E) Optical section through the tentacle bases (10-tentacle larva). (F1–F4) Telotroch: (F1, F3) general (F1) lateral view and (F3) viewed from the bottom (F2, F4) posterior EdU⁺ pre-telotrochal and post-telotrochal zones in 10- (F2) and 12- tentacle larvae (F4). (A1–E2; F2, F4) CLSM; (F1, F3) SEM. Magenta – EdU-click; yellow – anti-phosphorylated histone H3 IHC; cyan – nuclei stained with DAPI. Abbreviations: A – anterior, ao – apical organ, c1 – protocoel, c3 – metacoel, ct – central ciliated zone of telotroch, D – dorsal, es – esophagus, m – mouth, md – midgut, ms - metasomal sac, of - oral field, P – posterior, pl - preoral lobe, poz - post-telotroch proliferation zones, pr – proctodaeum, prz - pre-telotroch proliferation zones, st – stomach, t – tentacle, tt – telotroch, V – ventral. Scale bars: A1–A3, F3 – 50 µm; B, C, F4 – 20 µm; D, F2 – 15 µm; E – 5 µm; F1 – 25 µm.

In the nascent metasomal sac, proliferating cells are evenly distributed along both the epithelial and coelomic linings (Fig. 3D1). A comparison with the 12-tentacle stage (Fig. 3D2) indicates that the sac grows uniformly without forming a defined proliferation zone, which suggests balanced and continuous expansion of both layers.

In the telotroch, two distinct proliferation domains are already apparent at the 10-tentacle stage: a pre-telotrochal and a post-telotrochal zones, each consisting of EdU⁺ and pH3⁺ cells arranged in circular patterns (Fig. 3D, F1–F4). Initially, additional mitotic cells are also found between these rings, but by the 12-tentacle stage, they are separated by a distinct band of differentiated ciliated cells. A few dividing cells can still be detected between the anus and the telotroch at the 10-tentacle stage, but these are almost completely absent by the 12-tentacle stage (Fig. 3F3, F4), indicating the onset of functional compartmentalization at the posterior pole.

As larval development proceeds, further compartmentalization of proliferation domains becomes evident (Fig. 4A1–C). In the preoral lobe, proliferating cells become concentrated dorsally, with a predominance of cell divisions in the marginal zone. The mouth region, pharynx, and esophagus remain active proliferation sites up to the pre-competent stage (Fig. 4D). EdU incorporation continues prominently in the epidermis beneath the tentacles, while proliferative activity in the trunk epithelium at the level of the stomach declines significantly (Fig. 4B, C).

**Figure 4.**
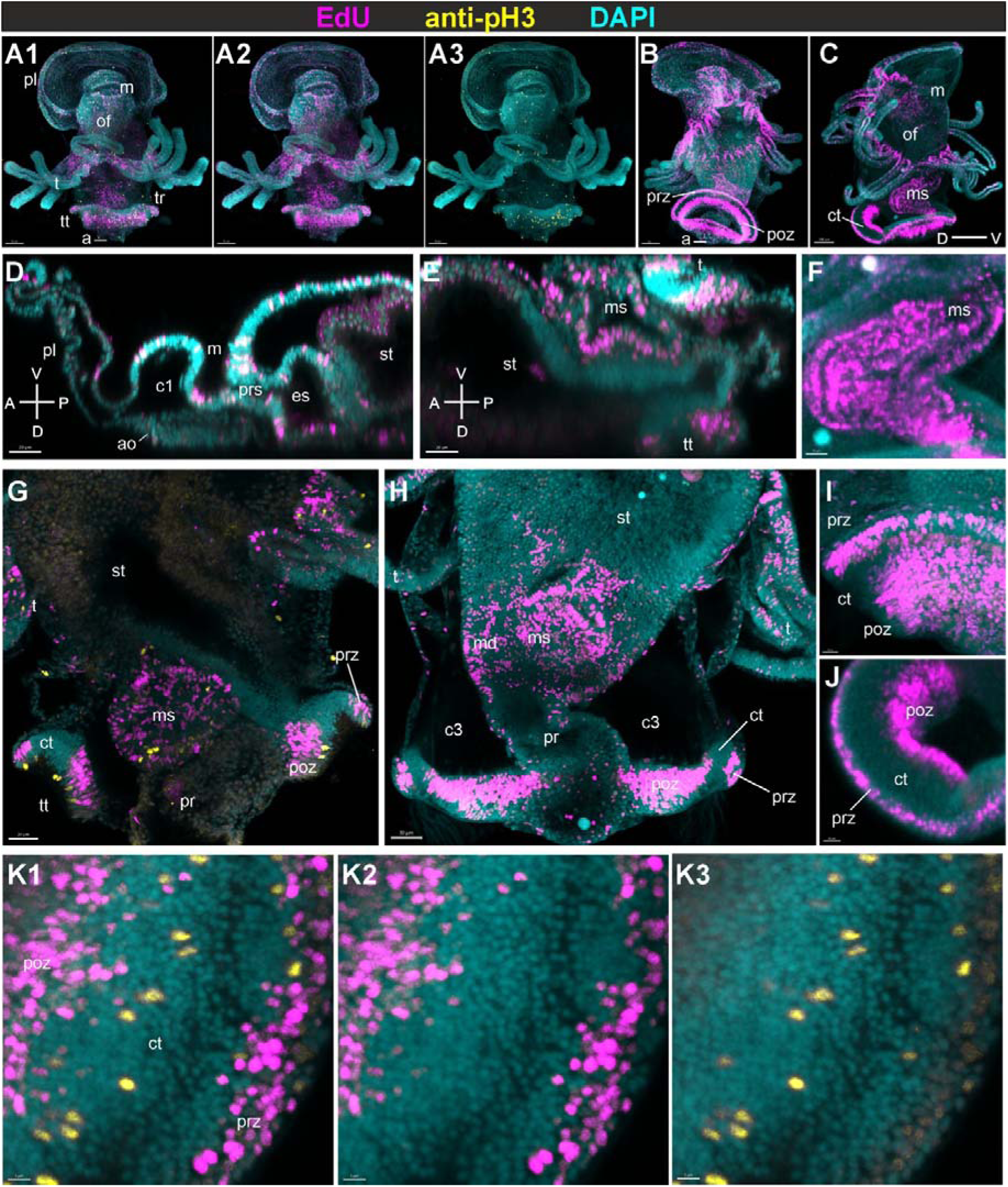
DNA replication and mitoses in mid and late larvae of *Phoronopsis harmeri*. (A1–C) Whole larvae: (A1–A3) 16-tentacle larva (ventral view); (B) 20-tentacle larva (dorsal view); (C) pre-competent larva (lateral view). (D, E) Sagittal sections of a 20-tentacle larva: (D) anterior region; (E) metasomal sac. (F) Sagittal section through the metasomal sac of a pre-competent larva. (G, H) Frontal sections of the metasomal sac in posterior regions of 16- (G) and 20- tentacle (H) larvae. (I–K3) Telotroch regions with EdU⁺ and pH3⁺ cells: (I) 16-tentacle larva; (J) pre-competent larva; (K1–K3) 20-tentacle larva. (A1–K3) CLSM. Magenta – EdU-click; yellow – anti-phosphorylated histone H3 IHC; cyan – nuclei stained with DAPI. Abbreviations: A – anterior, ao – apical organ, c1 – protocoel, c3 – metacoel, ct – central ciliated zone of telotroch, D – dorsal, es – esophagus, m – mouth, md – midgut, ms - metasomal sac, of - oral field, P – posterior, pl - preoral lobe, poz - post-telotroch proliferation zones, pr – proctodaeum, prs – prestomach, prz - pre-telotroch proliferation zones, st – stomach, t – tentacle, tt – telotroch, V – ventral. Scale bars: A1–A3 – 50 µm; B – 70 µm; C – 100 µm; D, E, G, J – 20 µm; F, H – 30 µm; I – 10 µm; K1–K3 – 5 µm.

In the tentacles, in addition to sustained proliferation at their bases, numerous mitotic cells are observed along their entire length, except at the distal tip on the anterior side (Fig. 4B, C). The entire wall of the elongating metasomal sac remains a site of intense EdU incorporation and mitotic activity (Fig. 4E–G). Within the gut, active cell division persists in the midgut and proctodeum, but not in the stomach (Fig. 4G, H). In the telotroch, both pre- and post-telotrochal proliferation zones remain distinct and active (Fig. 4G–K3), with the post-telotrochal domain showing a noticeable expansion in the number of EdU⁺ nuclei.

By the competent larva stage, the larval body reaches its maximum size, and the telotroch becomes fully developed and morphologically prominent (Fig. 5A1–B). Proliferative activity remains detectable but becomes more spatially restricted. In the preoral lobe, mitotic cells are concentrated primarily in the dorsal marginal zone, with scattered divisions in the central region (Fig. 5C1–D2). Optical sections reveal EdU⁺ nuclei in both the surface epithelium and deeper cell layers of this region (Fig. 5E).

**Figure 5.**
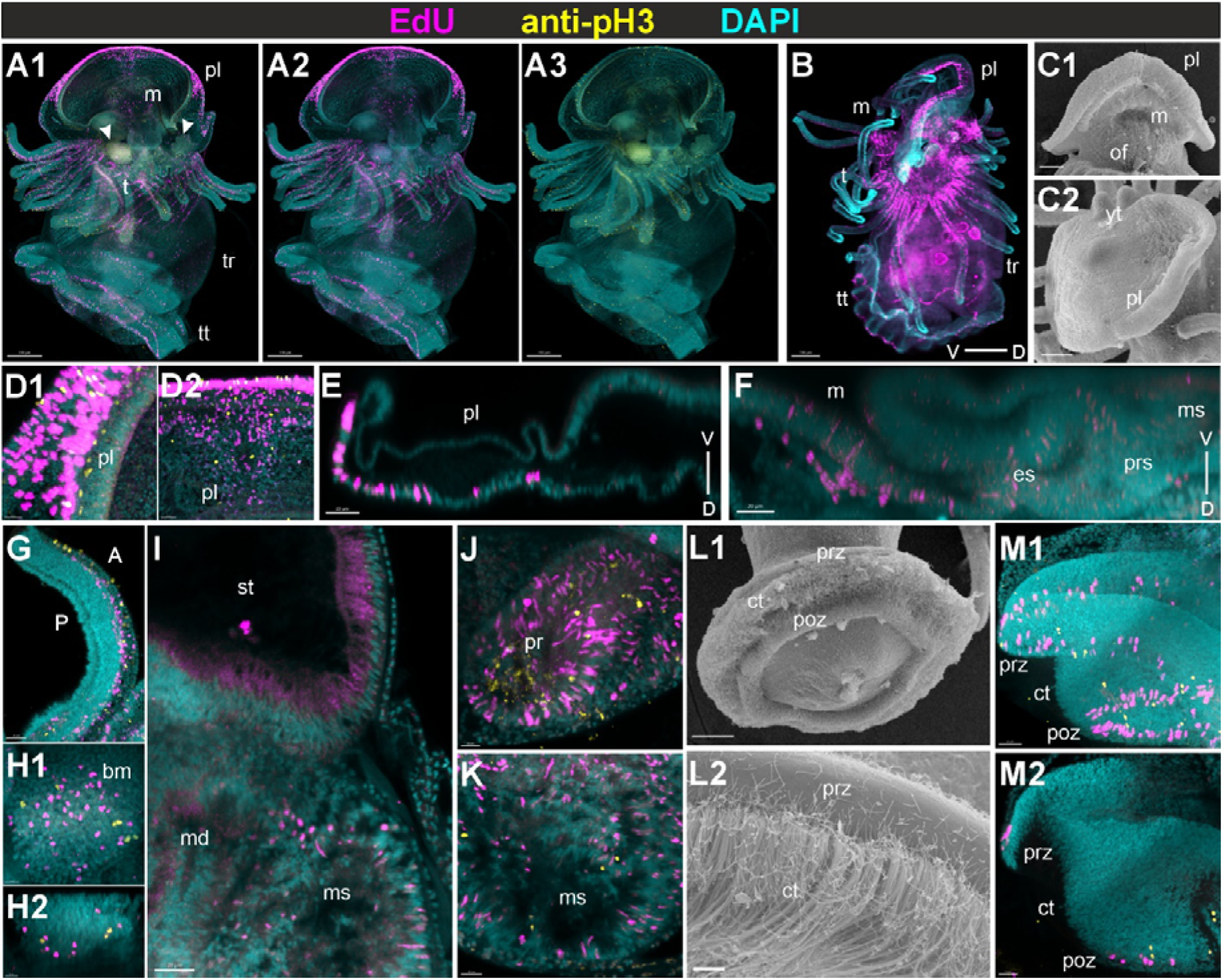
Proliferative activity in a competent larva of *Phoronopsis harmeri*. (A1–B) Whole larvae: (A1–A3) dorsal view; (B) lateral view. (C1, C2) The preoral lobe: (C1) ventral view; (C2) apical view. (D1, D2) Dividing cells in the preoral lobe: epithelium near the edge at (D1) the lateral and (D2) the central regions. (E, F) Sagittal optical sections through the preoral lobe (E) and mouth region (F). (G) Tentacle. (H1, H2) Blood mass (hemopoietic organ): (H1) frontal view; (H2) parasagittal section. (I) Frontal section through the stomach and midgut. Note the near absence of mitoses in both the stomach and adjacent epithelia. (J) Proctodeum. (K) Metasomal sac. (L1–M2) Telotroch: (L1) bottom view; (L2) bundles of cilia of the telotroch central zone; telotroch lateral region (M1) maximum intensity projection and (M2) frontal optical section. (A-B; D1-K; M1-M2) CLSM; (C1-C1; L1-L2) SEM. Magenta – EdU-click; yellow – anti-phosphorylated histone H3 IHC; cyan – nuclei stained with DAPI. Arrowheads – blood mass. Abbreviations: A – anterior, bm - blood mass, ct – central ciliated zone, of telotroch, D – dorsal, es – esophagus, m – mouth, md – midgut, ms - metasomal sac, of - oral field, P – posterior, pl - preoral lobe, poz - post-telotroch proliferation zones, pr – proctodaeum, prs – prestomach, prz - pre-telotroch proliferation zones, st – stomach, t – tentacle, tr – trunk, tt – telotroch, V – ventral, yt - youngest tentacle. Scale bars: A1–A3, B – 150 µm; C1–C2, L1 – 50 µm; D1, H1–H2, L2 – 10 µm; D2, J, M2 – 15 µm; E, F, G, I, K, M1 – 20 µm.

The mouth area and adjacent pharyngeal epithelium continue to show elevated proliferation (Fig. 5F), suggesting ongoing tissue remodeling during the final larval preparations for metamorphosis. Tentacle bases retain high mitotic activity, while additional divisions are observed along the proximal two-thirds of tentacle shafts (Fig. 5G), contrasting with earlier stages where divisions were mostly basal.

The stomach wall shows an almost complete absence of dividing cells (Fig. 5I), and this reduction in mitotic activity extends to adjacent tissues, including the coelomic lining and the overlying epidermis in the central region of the trunk. In contrast, the midgut and proctodeum continue to exhibit prominent EdU incorporation and mitoses, maintaining their status as active proliferation zones at this stage (Fig. 5J).

In the metasomal sac, EdU⁺ and pH3⁺ cells persist across the entire inner surface (Fig. 5K), indicating that this structure remains a site of robust cell proliferation up to metamorphic competence.

The blood masses (hemopoietic organ), which reaches the full size in competent larvae, shows localized proliferation confined to its deeper (ventral) regions (Fig. 5H1, H2), suggesting region-specific cell renewal or differentiation. In the telotroch, the pre- and post-telotrochal mitotic zones remain structurally defined and mitotically active (Fig. 5L1–M2), forming two concentric rings of proliferating cells encircling the posterior pole. Notably, the pre-telotrochal zone becomes less prominent than the post-telotrochal zone at this stage. The clear compartmentalization of these domains likely reflects the finalized spatial organization of larval structures prior to the onset of metamorphic remodeling.

At the onset of metamorphosis, proliferative activity becomes spatially reorganized in accordance with the transition from larval to juvenile body architecture. In early metamorphic individuals, EdU⁺ nuclei are still present in the regions inherited from the larval stage but become progressively restricted to domains contributing to juvenile structures (Fig. 6A1–A3). By the mid-metamorphic stage, proliferative activity becomes more compartmentalized and correlates with sites of morphogenetic remodeling (Fig. 6B1–B3).

**Figure 6.**
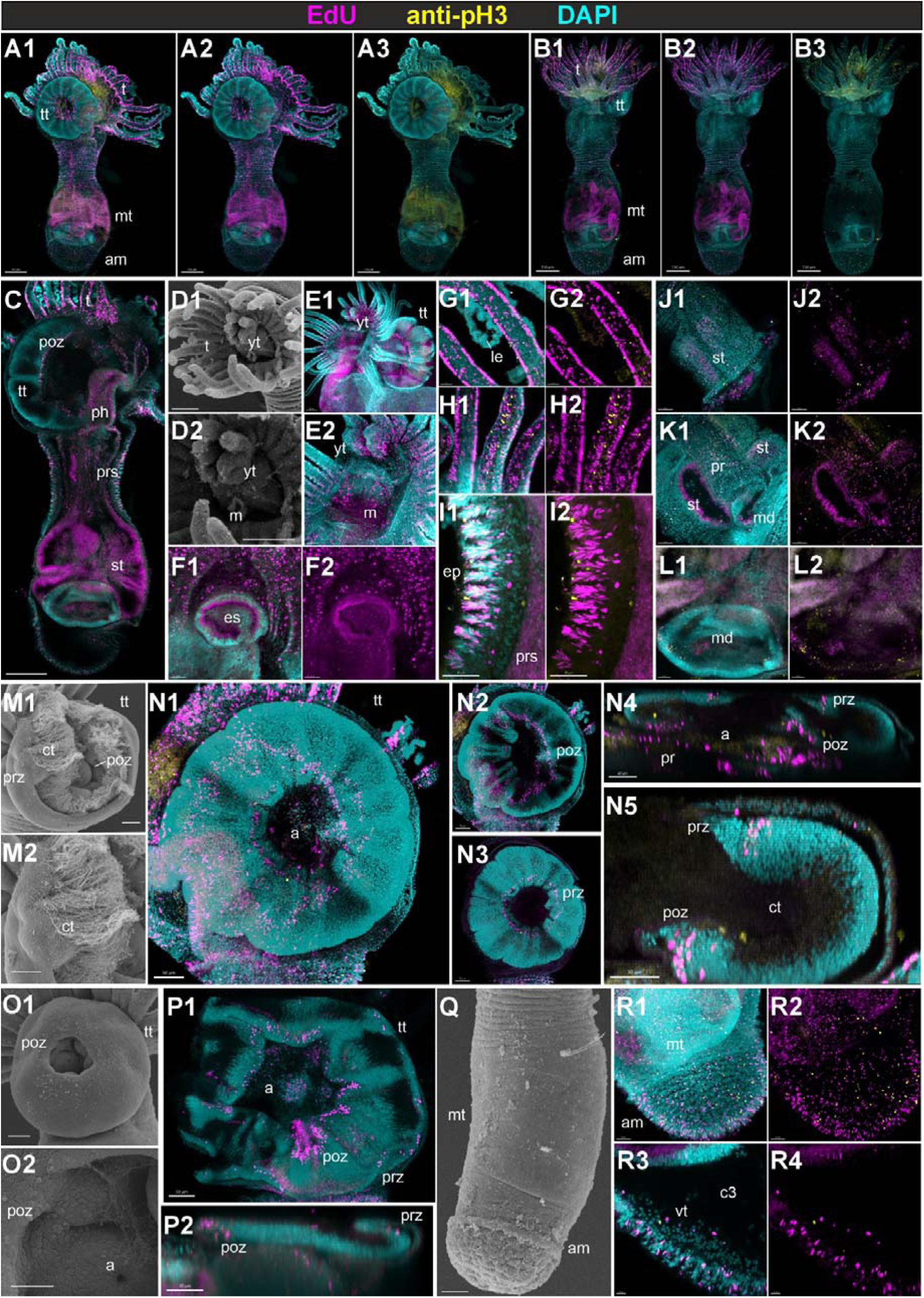
Cell proliferation during metamorphosis of *Phoronopsis harmeri*. (A1–B3) Whole juveniles: (A1–A3) early metamorphic stage (posterior view); (B1–B3) mid metamorphic stage (anterior view). (C) Frontal section at the level of the esophagus in the early metamorphic juvenile. (D1, D2) The mouth region in the mid metamorphic juvenile. (E1, E2) EdU⁺ nuclei in the mouth region (maximum intensity projection). (F1, F2) Frontal sections through the esophagus. (G1–H2) Proliferating cells in tentacles: (G1, G2) early stage; (H1, H2) mid stage. (I1, I2) Epidermal divisions near the head. (J1, J2) Epidermal divisions at the level of the stomach. Note the reduced number of mitoses in comparison with esophagus and midgut (I1-I2). (K1, K2) Frontal sections at the level of the stomach and proctodeum. (L1, L2) Frontal sections at the midgut level. Note fewer mitotic cells in the stomach wall compared to midgut and proctodeum. (M1–P2) Telotroch of metamorphic animal: (M1–N5) early stage; (O1–P2) mid stage. (N1) Remnant of the telotroch in posterior projection; (N2, N3) pre-telotrocgal and post-telotrochal proliferation zones; (N4, N5) sagittal sections. (P1, P2) The telotroch remnant in mid-metamorphic animal (P1) posterior view and (P2) sagittal section through the telotroch. (Q–R4) Ampulla of mid metamorphic animal: (Q) juvenile trunk and ampulla general view; (R1–R4) proliferation in the terminal region of the ampulla (R1, R2) maximum intensity projection) and (R3, R4) optical sections. Note the presence of mitotic cells scattered within the cavity. (A1-L2; N1-N5; P1-P2; R1-R4) CLSM; (M1-M2; O1-O2; Q) SEM. Magenta – EdU-click; yellow – anti-phosphorylated histone H3 IHC; cyan – nuclei stained with DAPI. Abbreviations: a – anus, am – ampulla, c3 – metacoel, ct – central ciliated zone of telotroch, ep – epithelium, es – esophagus, le - – sloughing off of the larval tentacle epidermis, m – mouth, md – midgut, mt – mid-trunk, ph – pharynx, poz – post-telotroch proliferation zone, pr – proctodaeum, prs – prestomach, prz - pre-telotroch proliferation zones, st – stomach, t – tentacle, tt – telotroch, vt – vasoperitoneal tissue, yt – youngest tentacle. Scale bars: A1–A3, B1–B3, C – 150 µm; E1 – 100 µm; E2, M1–M2, N1–N3, O1, P1, Q – 50 µm; G1–G2, H1–H2, O2, R1–R2 – 20 µm; I1–I2, L1–L2, N5, P2 – 30 µm; J1–J2, K1– K2, N4 – 40 µm; R3–R4 – 10 µm.

Proliferation persists in the esophageal region, which remains one of the most mitotically active parts of the digestive tract during metamorphosis (Fig. 6C). The mouth and pharyngeal areas also retain a high density of EdU⁺ nuclei, both in surface epithelia and in deeper layers of the oral cavity, reflecting continued growth and tissue renewal (Fig. 6D–F).

In the tentacles, superficial epithelial cells along the anterior face begin to slough off during early metamorphosis, and no dividing cells are detected in this layer. Instead, cell divisions are concentrated in the marginal zones of the tentacles. As metamorphosis progresses, proliferation becomes more prominent in the central core of the tentacles, suggesting a shift in proliferative focus toward internal structural reorganization (Fig. 6G1–H2).

The definitive epidermis of the juvenile trunk derives from the everted wall of the metasomal sac, where proliferative activity becomes compartmentalized already at the competent larva stage. This early regionalization is further elaborated during metamorphosis. In the epidermis near the anterior body region, adjacent to the forming head, proliferating cells exhibit a structured distribution: EdU⁺ nuclei are concentrated in the ridges of the developing surface, while the intervening grooves contain markedly fewer dividing cells, producing a patterned texture (Fig. 6I1, I2). In contrast, the epidermis at the level of the stomach shows only sparse and evenly distributed mitoses, with no evidence of ridge-groove organization (Fig. 6J1, J2).

The internal organs also maintain stage-specific proliferative patterns. The stomach epithelium continues to lack large detectable mitotic activity, consistent with its quiescent state since the competent larva stage. In contrast, the midgut and proctodeum remain proliferative, with EdU⁺ cells clearly visible in both regions (Fig. 6K1–L2), indicating ongoing epithelial turnover or reorganization.

In the telotroch, pre- and post-telotrochal proliferation zones persist into early metamorphosis. However, the pre-telotrochal domain becomes noticeably reduced and fragmented, while the post-telotrochal domain expands and almost merges with actively dividing cells near the anal opening, forming a broader proliferation zone at the posterior pole (Fig. 6M1–N5). As metamorphosis advances, the total number of dividing cells in the telotroch gradually declines, reflecting its ongoing resorption (Fig. 6O1–P2).

In the ampulla, a terminal, bulbous extension of the metasomal sac that gives rise to the anal region of the juvenile, EdU⁺ cells are distributed across the epidermis in a pattern resembling that of the anterior trunk. Proliferation is elevated in the forming ridges and reduced in the grooves, suggesting that this structure follows the same regionalization logic as the definitive trunk surface (Fig. 6Q–R4). Additionally, during mid-metamorphic stages, dividing cells are also observed within the internal cavity of the ampulla (Fig. 6R3, R4), indicating proliferative activity not only in the surface epithelium but also among dispersed cells within the coelomic cavity.

Together, these observations demonstrate that cell proliferation in *P. harmeri* larvae and juviniles is initially widespread but gradually becomes spatially organized. As development proceeds, distinct proliferation domains form in ectodermal, mesodermal and endodermal tissues, indicating a transition from generalized to patterned growth.

## Discussion

Our results reveal that during larval development of *Phoronopsis harmeri*, cell proliferation shifts from a broadly distributed pattern to a more regionally organized one as metamorphosis approaches. Mitotic activity becomes concentrated in key areas, including the mouth, esophagus, tentacle bases, metasomal sac, and especially the telotroch, where structured rings of dividing cells persist into late larval stages and metamorphic juveniles. We found that mitosis in this species displays several unconventional features, such as atypical chromatin organization and the absence of well-defined mitotic spindles. The spatial clarity and persistence of proliferation domains suggest a regulated and potentially lineage-specific mode of growth. Together, these observations indicate a high degree of spatial and cellular control over division, even in larvae with relatively simple body plans.

### Growth patterns and regionalization of proliferative activity in *Phoronopsis harmeri*

Our observations reveal several distinct features in the distribution and dynamics of cell proliferation during larval development and metamorphosis in *P. harmeri*.

As larval development proceeds, *P. harmeri* undergoes substantial body elongation. Notably, during the transition from the advanced actinotrocha to the competent larval stage (when elongation rates peak) cell divisions within the trunk become highly unevenly distributed. The central portion of the trunk shows minimal proliferative activity, while a prominent pre-telotrochal proliferation zone of dividing cells emerges at the boundary with the telotroch. This spatial pattern suggests that the pre-telotrochal domain likely contributes significantly to the elongation of the larval trunk. By mid-metamorphosis, however, mitotic activity in this ring markedly declines, whereas the post-telotrochal zone remains active. These observations imply that, beyond contributing to telotroch growth, the two proliferative zones may fulfill distinct developmental roles: the pre-telotrochal zone likely drives trunk elongation during larval life, whereas the post-telotrochal domain may be involved in the formation of the anal chamber of the juvenile gut, where *Cdx* expression is observed during metamorphosis (Temereva, 2010; Temereva and Kostyuchenko, 2025).

Importantly, *P. harmeri* larvae undergo not only linear body elongation but also substantial changes in body proportions. Notably, the telotroch expands disproportionately relative to total body size, while the preoral lobe maintains growth that is more proportional. The telotroch, composed of monociliated cells, serves as the primary locomotor organ in the planktonic larva. In phoronid species with relatively large increase of larval body size, such as *P. harmeri*, efficient ciliary locomotion may require increased cilia density. This, in turn, necessitates spatial expansion of the telotroch field, which appears to be achieved through proliferative mechanisms specifically localized to the telotroch epithelium. These divisions are spatially organized into pre- and post-telotrochal zones, which remain active throughout larval growth and into early metamorphosis, suggesting a mechanism that enables continued telotroch function during its expansion.

The tentacles originate from a broad and continuous zone of proliferation along the tentacle ridge, as previously described (Temereva and Kostyuchenko, 2025). As they grow, proliferative activity becomes increasingly restricted to the lateral bases of the tentacles, corresponding to the postoral ciliated band. This spatial restriction likely reflects a shift from initial patterning to maintenance and elongation, and parallels the proliferative patterns observed in other lophophorates such as bryozoans (Shunatova and Borisenko, 2020). Interestingly, the entire lophophoral region involved in tentacle formation displays sustained and spatially organized proliferative activity, and can therefore be regarded as a compartmentalized morphogenetic zone. It integrates spatially distinct subdomains that sequentially support tentacle patterning, initiation, and elongation. Such organization is reminiscent of growth zones in other radial or looped feeding structures among lophotrochozoans (Nielsen, 1994). This suggests that the lophophore functions not only as a definitive feeding apparatus, but also as a dynamic proliferative field actively contributing to morphological elaboration during both larval and postmetamorphic development.

The epithelium of the preoral lobe edge exhibits sustained proliferative activity until the onset of metamorphosis. This observation is particularly intriguing, given that the preoral lobe including the distal margin is entirely resorbed during metamorphosis and does not contribute to any definitive juvenile structures in *P. harmeri* (Temereva and Malakhov, 2015). From an evolutionary perspective, this region corresponds to the preoral portion of the ancestral tentacle apparatus – likely present in the last common bilaterian ancestor (Malakhov and Gantsevich, 2022), but secondarily lost in all extant lophophorates (Temereva, 2025). The persistence of proliferation in this transient, non-retained domain may represent a form of developmental atavism, reflecting residual activity of an evolutionarily suppressed morphogenetic program. Alternatively, this proliferative activity may serve a temporary functional role during the complex metamorphic transformation. *P. harmeri* larvae perform active, coordinated movements during metamorphosis. Although the edge of the preoral lobe does not exhibit overt morphological transformations in competent larvae, it may still play a mechanical or structural role in these morphogenetic events, for example, by locally modifying edge morphology and features. In addition to this potential role in metamorphosis, the preoral lobe serves several important functions during larval life. It bears ciliated and sensory cells involved in mechanoreception and chemoreception, likely contributing to spatial orientation, environmental scanning, and settlement behavior in the planktonic phase (Temereva and Wanninger, 2012; Temereva and Tsitrin, 2014). While the preoral lobe does not perform direct particle capture like the tentacles, it contributes to feeding by modulating ciliary currents and directing food-bearing water toward the mouth, acting in concert with the telotroch and tentacle crown (Temereva and Tsitrin, 2014). It may aid in hydrodynamic stabilization and maneuverability by shaping water flow in conjunction with the telotroch – a role suggested by flow modeling and structural observations (Johnson, 1988). Together, these sensory and biomechanical functions underscore the larval relevance of the preoral lobe, even if its distal edge is evolutionarily reduced and ultimately resorbed during metamorphosis.

The gut also shows a dynamic pattern of proliferation. In early larvae, both anterior and posterior regions of the gut actively divide, whereas the stomach proper exhibits few EdU⁺ or pH3⁺ cells. The growth of the stomach appears to occur primarily through proliferation at its anterior and posterior boundaries, possibly combined with local cell rearrangements. These proliferative dynamics are consistent with the known morphogenetic transformation of the larval gut into the definitive digestive tract during metamorphosis. In particular, the definitive prestomach develops from the upper portion of the larval stomach, which undergoes dramatic elongation and thinning, while the lower portion of the larval stomach becomes the juvenile stomach proper (Temereva, 2010). This remodeling corresponds well with the observed regionalization of proliferative zones and supports the notion that localized proliferation is tightly coupled with the spatial reorganization of the gut (Temereva and Kostyuchenko, 2025).

Another morphogenetically important domain is the coelomic cavity. At late larval stages, the mesothelial lining adjacent to the stomach exhibits minimal proliferative activity – a pattern mirrored in the neighboring ectodermal epithelium and coelomic lining. This reduction in proliferation is mirrored in the adjacent ectodermal epithelium and the coelomic lining in both competent larvae and postmetamorphic juveniles, where these epithelial surfaces are replaced by the everted wall of the metasomal sac. These tissues appear to enter a quiescent state, suggesting coordinated downregulation of proliferation across germ layers. Such regulation may reflect conserve d interlayer signaling, as documented in mollusks, annelids, and vertebrates, where mesodermal cues (e.g., FGF, TGF-β) shape ectodermal and endodermal development (Bennett, 1973; Grapin-Botton and Melton, 2000; Niu et al., 2010). Alternatively, the differentiated larval stomach may function as a local signaling center that inhibits proliferation in adjacent tissues, as described for mature endodermal domains in other systems (Horb and Slack, 2001; Okegbe and DiNardo, 2011; Zhu et al., 1999).

In contrast to the quiescent mesothelial domain near the stomach, proliferative activity is consistently observed in the coelomic lining of the ampulla during metamorphosis that gives rise to the juvenile vasculature and vasoperitoneal tissue. Ultrastructural studies indicate that the ampullar coelomic lining is composed of flattened mesothelial cells closely associated with underlying extracellular matrix components and embedded myofilaments, which likely contribute to vasomotor control and coelomic fluid regulation (Temereva et al., 2001). During metamorphosis, this lining plays a pivotal role in forming the definitive blood vessel walls and supporting tissues of the juvenile circulatory system, including the hemal sinuses and connective elements around the gut and ampulla Since the ampullar coelomic lining gives rise to the juvenile vasculature and associated vasoperitoneal tissues, the observed presence of EdU⁺ and pH3⁺ cells in this region likely reflects ongoing proliferation-driven differentiation processes. This is consistent with the broader pattern observed in phoronid development, where mesodermal proliferation remains active in specific regions contributing to organogenesis, despite the overall decline in mitotic activity in many trunk compartments at late larval stages (Temereva, 2024).

Ultrastructural studies have shown that the epithelial folds of the juvenile trunk in phoronids contain a dense network of basal muscle fibers and well-developed apical junctional complexes (Temereva, 2024; Temereva et al., 2001). These folds are not merely surface undulations but represent deep epithelial invaginations underlain by bundles of myoepithelial cells and rich innervation. The alternation of folds and grooves thus reflects a complex morphofunctional compartmentalization of the body wall. The localization of EdU⁺ nuclei within the folds suggests that these regions may serve as active growth zones required for the continued expansion or remodeling of the contractile epidermis. Given the biomechanical demands on the body wall during the early benthic life of phoronids, proliferative activity in folds may support not only epithelial renewal but also the reinforcement or differentiation of muscle components within the epithelial-muscular sac (Temereva, 2024; Temereva and Malakhov, 2015).

The mid-trunk region, corresponding anatomically to the stomach level, also displays distinctive histological and ultrastructural features. Its epidermis consists of hypertrophic secretory cells rich in electron-dense granules, likely involved in tube formation or protective coating (Temereva et al., 2001). Underlying these is a thick basal layer of circular muscle fibers forming a prominent myoepithelial system, composed of large, metabolically active contractile cells packed with actin-myosin filaments, mitochondria, and smooth endoplasmic reticulum. These features suggest that growth in this region may proceed through cellular hypertrophy rather than local proliferation. Additionally, expansion may be supported by the rearrangement and epithelial integration of proliferative cells originating from adjacent regions with higher mitotic activity. This combination of hypertrophy and targeted cell recruitment likely supports the biomechanical demands and compartmentalization of the mid-trunk body wall as it transitions from morphogenesis to functional specialization.

### Phoronid growth reveals coexistence of both conserved and novel proliferative mechanisms

The distribution of proliferation domains in *P. harmeri* larvae provides important insights into the evolutionary plasticity of growth strategies among lophotrochozoans. While many bilaterians rely on localized zones of proliferation – so-called posterior growth zones (PGZ) – for axial elongation and segment formation, the patterns observed in *P. harmeri* suggest a more diverse and modular approach to body growth. This makes phoronids an informative group for comparing conserved developmental mechanisms with lineage-specific innovations.

The identification of PGZs in annelids and panarthropods (including arthropods, onychophorans, and tardigrades) has played a central role in the debate on the homology of segmentation across Bilateria, with many studies proposing a shared segmentation mechanism driven by a PGZ (Balavoine and Adoutte, 2003; Scholtz, 2002). However, the distribution of proliferation domains in *P. harmeri* underscores the evolutionary diversity of growth strategies within Lophotrochozoa.

In annelids, the PGZ consists of a localized pool of proliferating stem-like cells that contribute to segment formation in a sequential manner. Studies on polychaetes like *Platynereis dumerilii* and *Capitella teleta* reveal that this process is governed by a conserved molecular network, with Wnt/β-catenin, FGF, and Notch signaling coordinating the posterior proliferation and patterning of new segments (De Rosa et al., 2005; Seaver et al., 2005). This proliferative region ensures a continuous supply of undifferentiated cells for segment addition, a feature that is also observed in clitellates, where teloblastic stem cells undergo highly ordered divisions to generate new segmental structures (Matsuo et al., 2005; Rivera et al., 2005). However, while this localized proliferative mechanism is well-documented in annelids, the situation in panarthropods is complex. While arthropods generally display posterior growth-driven segmentation, direct evidence for a PGZ is limited to certain crustaceans such as malacostracans, where a posterior pool of stem-like cells has been identified (Dohle and Scholtz, 1988; Scholtz, 2002). In contrast, in most arthropods, segmentation relies on a molecular segmentation clock, where oscillatory gene expression regulates segment addition rather than a strict PGZ-driven process (Chipman, 2010; Liu and Kaufman, 2005). This divergence raises significant questions about the ancestral segmentation mechanism in panarthropods and whether annelid and arthropod segmentation share a deep homology. Adding to this debate, studies on onychophorans (velvet worms) suggest that they lack a localized PGZ entirely, challenging previous assumptions that a posterior proliferation zone was an ancestral trait of panarthropods (Mayer et al., 2010). Onychophoran embryos instead exhibit a distributed proliferation pattern, in which cell divisions occur more evenly along the elongating body axis rather than being restricted to a posterior zone. This finding undermines the idea that a proliferative PGZ is a universal feature of segmented bilaterians and supports the hypothesis that segmentation may have evolved independently in annelids and arthropods through modifications of an ancestral posterior elongation system (Jaeger and Goodwin, 2002). The absence of a posterior proliferation zone in onychophorans thus necessitates a reassessment of segmentation homology and suggests that the PGZ-driven segmentation mechanism observed in annelids may represent an independent evolutionary innovation rather than a shared ancestral feature of annelids and arthropods. Given these uncertainties, further comparative studies on cell proliferation patterns in other groups could provide critical insights into the evolutionary origins of segmentation and the role of posterior growth mechanisms in bilaterian development.

While annelids rely on a posterior proliferation zone, many non-segmented spiralians exhibit distinct growth strategies. Nemerteans, for example, lack a clearly defined PGZ but rely on localized cell proliferation zones to drive body elongation. The development of nemertean larvae, particularly pilidium larvae, involves a stem-cell-driven growth process that enables indirect development (Maslakova, 2010). These larvae harbor undifferentiated cells within the larval body, which later migrate and contribute to adult tissues (Bird et al., 2014). Similarly, mollusks, including bivalves and gastropods, exhibit unique patterns of cell division. Studies on *Pinctada fucata* (a pearl oyster) have demonstrated that mantle tissue growth relies on the proliferation of epithelial and mesenchymal cells in specific marginal and central zones (Fang et al., 2008). In gastropods, on the one hand, foot elongation occurs through scattered cell divisions without the involvement of PGZ (Glebov et al., 2014; Kurtova et al., 2024). On the other hand, the torsion process is driven by differential cell proliferation on opposite sides of the body (Kurita and Wada, 2011), showcasing evolutionary innovations rather than conserved developmental mechanisms. Interestingly, unlike the homologous foot of gastropods, the elongation of cephalopod arms is associated with the formation of distal proliferation zones, which may have evolved secondarily (Nödl et al., 2016, 2015). This suggests a fundamental shift in the spatial organization of growth regulation within mollusks, where cephalopods developed a localized distal growth mechanism, contrasting with the distributed cell proliferation observed in gastropods. Overall, in contrast to annelids, molluscan development is highly regionalized, with proliferation occurring in specific domains rather than a continuous posterior zone. This highlights the evolutionary plasticity of morphogenetic processes in mollusks, where major body transformations can arise from localized shifts in cell division patterns rather than relying on ancestral growth strategies observed in other lophotrochozoans.

Platyhelminths (flatworms), which are acoelomate spiralians, exhibit even more divergent developmental patterns. Many free-living flatworms such as *Macrostomum lignano* rely on neoblasts – totipotent stem cells distributed throughout the body – for growth and regeneration, rather than a posterior proliferation zone (Egger et al., 2006). In flatworms that lack neoblasts, regeneration still occurs through scattered cell divisions within the corresponding tissues (Gąsiorowski et al., 2025). However, some parasitic platyhelminths exhibit localized posterior proliferation, particularly in their larval stages, where new body units such as proglottids are continuously generated in cestodes (Koziol et al., 2010). The presence or absence of a PGZ across lophotrochozoans raises fundamental questions about the evolution of body axis elongation and segmentation. The findings from annelids and their segmented PGZ-driven growth contrast with the non-segmented, regionally proliferative growth observed in mollusks, nemerteans, bryozoans, and platyhelminths. This suggests that posterior growth mechanisms have been independently modified across different lophotrochozoan lineages.

The presence of clearly defined proliferative zones in the telotroch and the persistence of their activity until metamorphosis suggests the possible existence of a PGZ homolog in the actinotroch larva of *P. harmeri*. Unlike in annelids, where the PGZ contributes to segment formation, this proliferation zone likely supports larval axial elongation and maintenance of the posterior epithelium without generating repeated morphological units. Such a PGZ may represent either a conserved bilaterian feature or a secondary adaptation linked to the larval mode of development. At the same time, as in nemerteans and gastropods, proliferative domains in *P. harmeri* are not restricted to a posterior zone but are distributed across multiple organs and body regions. For example, both the metasomal sac and juvenile trunk epidermis exhibit widespread mitotic activity without a clearly localized growth center, suggesting a mode of regional rather than axial growth. This pattern contrasts with the strategies observed in many other spiralians discussed above, where either localized PGZs or stem-cell-based systems dominate axis elongation and tissue formation. Instead, *P. harmeri* displays a hybrid proliferative strategy that combines sustained posterior growth with regionally organized proliferation, reflecting a distinct developmental mode that may have evolved in response to its specific life history and morphological constraints. The combination of both possibly conserved (e.g., telotrochal PGZ) and novel (e.g., regionally organized metasomal proliferation) growth mechanisms highlights *P. harmeri* as a valuable model for reconstructing the evolution of growth strategies in Lophotrochozoa and for understanding the plasticity of axial and regional proliferative systems.

### Atypical aspects of mitosis and evidence for interkinetic nuclear migration in *Phoronopsis harmeri*

Certain tissues in multicellular animals exhibit naturally modified mitotic processes. For example, mammalian oocytes are known to undergo acentriolar, spindle-deficient divisions where chromosomes are segregated without the involvement of canonical centrosomes (Dziugiel, 2014). Similarly, in early marine invertebrate embryos, such as echinoderms and cnidarians, cell divisions sometimes rely on modified spindle structures or unorthodox metaphase configurations (McIntosh and Hays, 2016). Nonetheless, these are typically transient developmental adaptations rather than fixed features of adult somatic mitosis. The situation in *P. harmeri*, where this atypical mitosis is observed in differentiated epithelial tissue, indicates a fundamentally different pattern, where the non-canonical division mode may be intrinsic rather than transient.

A broad spectrum of atypical mitotic features is observed among unicellular eukaryotes, often as a result of specialized evolutionary pressures. In *Trypanosoma*, for example, cell division proceeds via a unique mechanism in which chromosomes do not align on a metaphase plate and kinetochores are poorly defined or absent (Akiyoshi and Gull, 2013; Smith, 2013). Although mechanistically distinct, the disorganized metaphase and absence of visible spindle structures in *P. harmeri* bear a conceptual resemblance to these protist strategies. However, whether the ultrastructural features of mitotic nuclei observed by TEM in phoronids truly reflect the absence of specific canonical components involved in chromosome segregation and cell division remains to be determined. It is particularly remarkable that such a system is maintained in a multicellular context and in differentiated epithelial tissues, raising important questions about the evolutionary plasticity of mitotic architecture in metazoans.

Another notable feature of mitosis in *P. harmeri* is the spatial separation of proliferative phases both within the pseudostratified telotroch epithelium and in the larval trunk epidermis. EdU⁺ nuclei, marking cells in S-phase, are consistently localized near the basal surface, whereas pH3⁺ mitotic figures cluster apically. This apicobasal segregation suggests the presence of interkinetic nuclear migration (INM), a process best characterized in vertebrate neuroepithelia, where the nuclei of cycling cells migrate along the apicobasal axis in synchrony with the cell cycle (Kosodo et al., 2011). In such systems, DNA synthesis (S-phase) occurs basally and mitosis apically, maintaining orderly proliferation and epithelial integrity during development. INM plays a key role in the formation of the complex multilayered architecture of the vertebrate’s brain (Laguesse et al., 2015). INM-like processes have also been reported in different invertebrate groups. Signs of INM have been observed in the cnidarian *Nematostella vectensis* (Meyer et al., 2011; Nakanishi et al., 2012), and nuclear movements along the apicobasal axis have been described during echinoderm development (Mercurio et al., 2024). In *Drosophila*, pseudostratified epithelial tissues such as imaginal discs and the eye primordium show regulated nuclear positioning along the apicobasal axis, driven by actomyosin and microtubule-based mechanisms (Mosley-Bishop et al., 1999; Badugu and Käch, 2020). Among lophotrochozoans, evidence of INM has been reported in the neurodevelopment of annelid *Platynereis* (Tosches, 2013), supporting the view that INM represents a fundamental and conserved mechanism in pseudostratified epithelia of multicellular animals (Meyer et al., 2011). In phoronid, we observed this phenomenon in epithelial tissues not clearly associated with neurogenesis and it appears to be broadly involved in larval development. Interestingly, signs of INM are found not only in the pseudostratified telotroch epithelium but also in the monolayered trunk epidermis. Overall, our observations in phoronid may be of particular interest for future studies on conserved and convergent mechanisms of intracellular nuclear dynamics.

## Conclusions

Our analysis of proliferative activity in *Phoronopsis harmeri* revealed a complex and dynamic landscape of larval and postmetamorphic growth. Rather than relying solely on a centralized posterior growth zone (PGZ), *P. harmeri* exhibits a modular arrangement of proliferation domains that are spatially and functionally specialized. This includes persistent telotrochal rings, proliferative zones at tentacle bases, preoral and postoral regions, and mesodermal compartments such as the metasomal sac and ampulla.

The coexistence of regionally organized epithelial proliferation and possible homologs of PGZs suggests that *P. harmeri* employs both conserved and lineage-specific strategies to support body elongation, organogenesis, and metamorphic remodeling. Comparisons with other lophotrochozoans, including annelids, mollusks, and nemerteans, indicate that phoronid growth reflects an evolutionary mosaic: it integrates ancestral mechanisms of axial extension with novel, regionally defined proliferative modules.

These findings position phoronids as valuable models for understanding how diverse growth architectures evolve within Bilateria. Ultimately, *P. harmeri* highlights the plasticity of metazoan growth strategies and the evolutionary potential of spatially regulated cell proliferation.

## Acknowledgements

The authors are grateful to Dr. Elena Voronezhskaya for her support and insightful discussions. This research was carried out using the facilities of the Core Centrum of the Institute of Developmental Biology RAS under IDB RAS RP No. 0088-2024-0015. This study was conducted with financial support from the Russian Science Foundation (#23-14-00020) and under the state assignment of Lomonosov Moscow State University. In this study, the scientific equipment, which was purchased in frame of special MSU program of development, has been used.

## Notes

### Competing Interest Statement

The authors have declared no competing interest.

